# Impact of age and sex on neuroinflammation following SARS-CoV-2 infection in a murine model

**DOI:** 10.1101/2023.08.11.552998

**Authors:** Venkatramana D. Krishna, Allison Chang, Holly Korthas, Susanna R. Var, Walter C. Low, Ling Li, Maxim C-J. Cheeran

## Abstract

Severe Acute Respiratory Syndrome Coronavirus 2 (SARS-CoV-2), the etiological agent for the worldwide COVID-19 pandemic, is known to infect people of all ages and both sexes. Senior populations have the greatest risk of severe disease, and sexual dimorphism in clinical outcomes has been reported in COVID-19. SARS-CoV-2 infection in humans can cause damage to multiple organ systems, including the brain. Neurological symptoms are widely observed in patients with COVID-19, with many survivors suffering from persistent neurological and cognitive impairment, potentially accelerating Alzheimer’s disease. The present study aims to investigate the impact of age and sex on the neuroinflammatory response to SARS-CoV-2 infection using a mouse model. Wild-type C57BL/6 mice were inoculated, by intranasal route, with SARS-CoV-2 lineage B.1.351 variant known to infect mice. Older animals and in particular males exhibited a significantly greater weight loss starting at 4 dpi. In addition, male animals exhibited higher viral RNA loads and higher titers of infectious virus in the lung, which was particularly evident in males at 16 months of age. Notably, no viral RNA was detected in the brains of infected mice, regardless of age or sex. Nevertheless, expression of IL-6, TNF-α, and CCL-2 in the lung and brain was increased with viral infection. An unbiased brain RNA-seq/transcriptomic analysis showed that SARS-CoV-2 infection caused significant changes in gene expression profiles in the brain, with innate immunity, defense response to virus, cerebravascular and neuronal functions, as the major molecular networks affected. The data presented in this study show that SARS-CoV-2 infection triggers a neuroinflammatory response despite the lack of detectable virus in the brain. Age and sex have a modifying effect on this pathogenic process. Aberrant activation of innate immune response, disruption of blood-brain barrier and endothelial cell integrity, and supression of neuronal activity and axonogenesis underlie the impact of SARS-CoV-2 infection on the brain. Understanding the role of these affected pathways in SARS-CoV-2 pathogenesis helps identify appropriate points of therapeutic interventions to alleviate neurological dysfunction observed during COVID-19.

## Introduction

Severe acute respiratory syndrome coronavirus 2 (SARS-CoV-2) is the causative agent of the coronavirus disease 2019 (COVID-19) pandemic. COVID-19 symptoms range from mild flu-like illness, fever, fatigue, dry cough and dyspnea to fatal pneumonia and acute respiratory distress (Huang et al., 2020). Since the beginning of the pandemic in March 2020, there have been more than 768 million confirmed cases of COVID-19, with almost 7 million reported deaths [1]. SARS-CoV-2 is an enveloped, single-stranded, positive sense RNA virus that belongs to the *Betacoronavirus* genus in the *Coronaviridae* family [2, 3]. SARS-CoV-2 consists of four structural proteins: spike (S), membrane (M), envelope (E), and nucleocapsid (N). The S protein mediates viral entry into host cells by binding to the surface receptor angiotensin-converting enzyme 2 (ACE2) [4].

Although SARS-CoV-2 primarily infects cells in the respiratory tract, it affects multiple organ systems, including the central nervous system (CNS) [5–7]. Nearly 50% of patients infected with SARS-CoV-2 experience a post-acute sequela after the initial infection known as long COVID [8]. Long COVID is characterized by numerous symptoms, including persistent fatigue and neurological issues, which may last for many months post-acute SARS-CoV-2 infection (Weinstock et al., 2021). In fact, at least 20-45% of individuals infected with SARS-CoV-2 experience an array of neurological and neurodegenerative issues and cognitive deficits (O’Mahoney et al., 2023; Patel et al., 2022; Cron et al., 2022; Merdaca et al., 2021). It has been hypothesized that the presence of viral antigens and chronic inflammation, including the activation of brain mast cells, microglia, astrocytes, and other immune cells, may influence the pathogeneis and duration of long COVID [9, 10].

Animal models play a crucial role in the study of COVID-19 pathogenesis. Wild type laboratory mice are not susceptible to initial lineage A variants of SARS-CoV-2 due to inefficient interaction between the S protein and mouse ACE2 receptor [11]. To overcome this limitation, various mouse models expressing human ACE2 (hACE2) have been utilized such as K18-hACE2 transgenic mice that express hACE2 under the control of the cytokeratin 18 promoter [12, 13], humanized ACE2 mice by replacing endogenous mouse ACE2 with hACE2 using CRISPR/Cas9 knockin technology [14], Ad5-hACE2 transduced mice [15], and AAV-hACE2 transduced mice [16]. Mouse-adapted strains of SARS-CoV-2 have also been developed by multiple laboratories that can infect wild type laboratory mice [17–20]. Further characterization of mouse-adapted SARS-CoV-2 by sequencing showed that the N501Y mutation in the receptor binding domain (RBD) of S protein increase virulence in mice [19]. Studies found that some of naturally occurring SARS-CoV-2 variants such as B.1.1.7, B.1.351, and P.1 variants also possess N501Y mutation and can efficiently infect wild type laboratory mice [21–26].

Clinical studies of patients with COVID-19 suggested that increased age is associated with severe outcome and higher mortality [27–30]. Hospital admissions and mortality rates were higher in patients over 65 years old [30]. In addition, epidemiological and clinical studies have shown that COVID-19 severity and mortality rates are higher in men than in women [31–34]. The overarching goal of this study is to assess the effect of age and sex on SARS-CoV-2 infection. We hypothesize that an increase in age is associated with enhanced neuroinflammation after SARS-CoV-2 infection, which impacts the severity of neurological outcomes in COVID-19. To test this hypothesis, the B.1.351 variant of SARS-CoV-2 was used to infect wild type C57BL/6J mice (male and female) at different ages to investigate the neuroinvasive potential of SARS-CoV-2 and associated neuroinflammation.

## Materials and Methods

### Cells and virus

Vero E6 cells (ATCC CRL-1586, Manassas, VA, USA) were grown in Dulbecco’s modified Eagle medium (DMEM) supplemented with 5% heat inactivated fetal bovine serum (FBS). SARS-CoV-2, isolate hCoV-19/South Africa/KRISP-EC-K005321/2020, NR-54008, lineage B.1.351 was obtained from BEI Resources, NIAID, NIH (Manassas, VA, USA) and propagated in Vero E6 cells. Authenticity of the viral strain was validated by genome sequencing and compared to pubished sequence (GISAID Accession ID EPI_ISL_678597). Virus titers were determined by focus-forming assay on Vero E6 cells. All procedures with infectious SARS-CoV-2 were performed in certified biosafety level 3 (BSL3) facilities at the UMN using appropriate standard operating procedures (SOPs) and protective equipment approved by the UMN Institutional Biosafety Committee.

### Mice

C57BL/6J mice (The Jackson laboratory Stock No. 000664) and K18-hACE2 mice (The Jackson laboratory, Stock No. 034860) were propagated at the UMN and maintained in specific pathogen-free conditions. Mice were housed in groups of 3 to 5 per cage and maintained on a 12 h light/12 h dark cycle with access to water and standard chow diet *ad libitum*. Both male and female C57BL/6J mice at different ages (4-, 10-, and 16-months) or 4-month-old hemizygous K18-hACE2 mice were used in this study.

### Infection of mice with SARS-CoV-2

Mice were randomly assigned to infected and uninfected groups with equal numbers of male and female mice in each group. Mice in the infected group were anesthetized with 3% isoflurane/1.5 L/min oxygen in an induction chamber and inoculated intranasally with 1 x 10^5^ focus forming unit (FFU) of SARS-CoV-2 in a volume of 50 μL DMEM, split equally between each nostril. Mice were monitored, and the body weight measured daily for the duration of the experiment. At 7 days post infection (dpi), mice were euthanized, and lungs and brains were harvested for downstream analysis.

### Viral RNA quantification by PCR

Approximately half of the lung and the left hemisphere of each brain were weighed and homogenized in 1 mL of DMEM supplemented with 2% FBS in GentleMACS™ M tube using GentleMACS™ Dissociator (Miltenyi Biotec) with RNA 2.01 program setting. Tissue homogenates were stored in aliquots at –80 °C until use. Total RNA was extracted from 250 μL of tissue homogenate using Trizol™ LS reagent (Thermo Fisher scientific, Waltham, MA) according to the manufacturer’s instructions. The purity of RNA was assessed by calculating the ratio of OD_260_/OD_280_. 1 μg of total RNA was reverse transcribed using High-Capacity cDNA reverse transcription kit (Thermo Fisher scientific, Waltham, MA) according to manufacturer’s instructions. RT-qPCR was performed using Fast SYBR Green master mix (Thermo Fisher scientific, Waltham, MA) in a 7500 Fast Real-time PCR system (Applied Biosystems) using the Centers for Disease Control and Prevention RT-PCR primer set targeting SARS-CoV-2 nucleocapsid (N) gene, forward primer 5’-GACCCCAAAATCAGCGAAAT-3’ (2019-nCoV_N1-F), reverse primer 5’-TCTGGTTACTGCCAGTTGAATCTG-3’ (2019-nCoV_N1-R). The following reaction conditions were used: 95 °C for 3 min followed by 40 cycles of 95 °C for 10 s, 65.5 °C for 30 s, and 72 °C for 30 s. A standard curve was generated to determine genome copy numbers in the tissue sample by performing a qRT-PCR using synthetic SARS-CoV-2 RNA control (Twist Bioscience, Cat #104043). Viral loads are presented as log_10_ copies per μg of total RNA.

### SARS-CoV-2 focus forming assay

Vero E6 cells were seeded in 24-well tissue culture plates at a density of 1 x 10^5^ cells/well and incubated until the monolayer was 90-100% confluent. Four replicate wells of Vero E6 cells were infected with 100 μL of 10-fold serial dilution of virus or lung homogenate and incubated for 1h with intermittent mixing at 37°C/ 5% CO_2_. After 1 h, 500 µl of overlay medium containing 1.6% microcrystalline cellulose, 2% heat-inactivated FBS in DMEM was added to each well and incubated for 48 h at 37°C/ 5% CO_2_. The cells were fixed with 4% paraformaldehyde for 30 min at room temperature. Fixed cells were washed with PBS containing 0.3% Triton X-100 (PBST), blocked with blocking buffer (1% bovine serum albumin (BSA) and 1% normal goat serum in PBST) for 1 h at room temperature, and then treated with rabbit anti-SARS-CoV-2 nucleocapsid antibody (Sino Biological; 1:2000 dilution) overnight at 4°C. After washing twice with PBST, cells were treated with alkaline phosphatase-conjugated goat anti-rabbit IgG (Thermo Fisher scientific, Waltham, MA; 1:1000 dilution) at room temperature for 1 h. Cells were washed twice with PBST and incubated for 20 min at room temperature in the dark with 1-Step™ NBT/BCIP substrate solution (Thermo Fisher scientific, Waltham, MA). After incubation, wells were washed with deionized water and the numbers of foci were counted. Infectious virus titer was expressed as focus forming units (FFU) per mL.

### Immunohistochemistry

Mouse brain hemispheres were drop-fixed with 4% paraformaldehyde (PFA) for 48 h and washed 3 times with 1X PBS. Brains were embedded in Tissue-Tek OCT (Andwin Scientific, IL) on dry ice. Frozen blocks were stored at –80°C until sectioning. Embedded brains were sagittally sectioned at 10 μm using a Cryostat (Leica Biosystems Inc). Sections were washed with wash buffer (0.3% Triton-X 100 in 1X PBS) and blocked with blocking buffer (1% normal goat serum, 0.3% Triton-X 100 in 1X PBS) for 60 min at room temperature. Tissue sections were incubated overnight at 4°C with primary antibody against the SARS-CoV-2 nucleocapsid protein (1:1000, Sino Biologicals US Inc, Wayne, PA). After washing 3 times with wash buffer, sections were incubated with goat anti-rabbit IgG-Alexa Fluor™ 594 (1:1000, ThermoFisher Scientific, Waltham, MA) for 60 min at room temperature. DAPI (4′,6-diamidino-2-phenylindole, ThermoFisher Scientific, Waltham, MA) was used to label nuclei. Images were captured using Leica DMi8 inverted microscope and Leica LAS software.

### RNA-Seq analysis

RNA was extracted from homogenized tissue as described above. Isolated RNA samples were submitted to the UMN Genomics Center core facility for library preparation and sequencing as previously described [35, 36]. Briefly, RNA quantities were determined by RiboGreen RNA Quantification assay (Thermo Fisher Scientific, Waltham, MA) and RNA quality was assessed using the Agilent Bioanalyzer (Agilent Technologies). Library preparation was performed using a TruSeq Stranded mRNA Library Prep Kit (Illumina) according to the manufacturer’s instructions. Sequencing was performed on a NovaSeq 6000 platform (Illumina), generating 20 million 150-bp paired-end reads per sample.

Raw sequencing reads were evaluated and trimmed using Trimmomatic (Bolger et al., 2014). Trimmed reads were mapped to the mouse reference genome GRCm38 using HISAT2 (https://doi.org/10.1038/nmeth.3317). RNA-seq raw counts for each gene were extracted using FeatureCounts [37]. Exploratory data analysis using PCA was performed on normalized data for outlier detection. One outlier (uninfected, male) was detected and excluded from subsequent analyses. Differential gene expression (DGE) analysis was conducted using DESeq2 (1.38.2) [38]. Default DESeq2 normalization and filtering methods were applied.

Data from both sexes (male and female) were analyzed in pairwise comparisons between infected and uninfected males and females, in addition to pairwise comparisons between infected and uninfected mice using the Wald test (design = ∼ sex + infection). Genes were considered significantly different between groups with an adjusted p value of < 0.05 after Benjamini-Hochberg correction. GO gene enrichment analysis was performed using the gseGO function in the ClusterProfiler package (4.6.0) [39]. Volcano plots were generated using ggplot2 (3.4.0). Hierarchical cluster analysis of the differentially expressed genes (DEGs) was performed using pheatmap (1.0.12) with default parameters (clustering_distance_cols = euclidean, clustering_method = complete). All DEG and pathway enrichment analyses were performed in R (4.2.1).

### Real-time quantitative polymerase chain reaction (RT-qPCR)

Total RNA was extracted from lung and brain homogenates and cDNA was synthesized from 1 μg total RNA as described above. The cDNA was amplified by RT-qPCR using Fast SYBR Green master mix (Thermo Fisher scientific, Waltham, MA) in a 7500 Fast Real-time PCR system (Applied Biosystems) using gene specific primers (**Table 1**). The specificity of RT-qPCR was assessed by analyzing the melting curves of PCR products. Ct values were normalized to RPL27 gene, and the fold change was determined by comparing virus-infected mice to uninfected controls using 2^-ΔΔCt^ method [40].

**Table 1.**
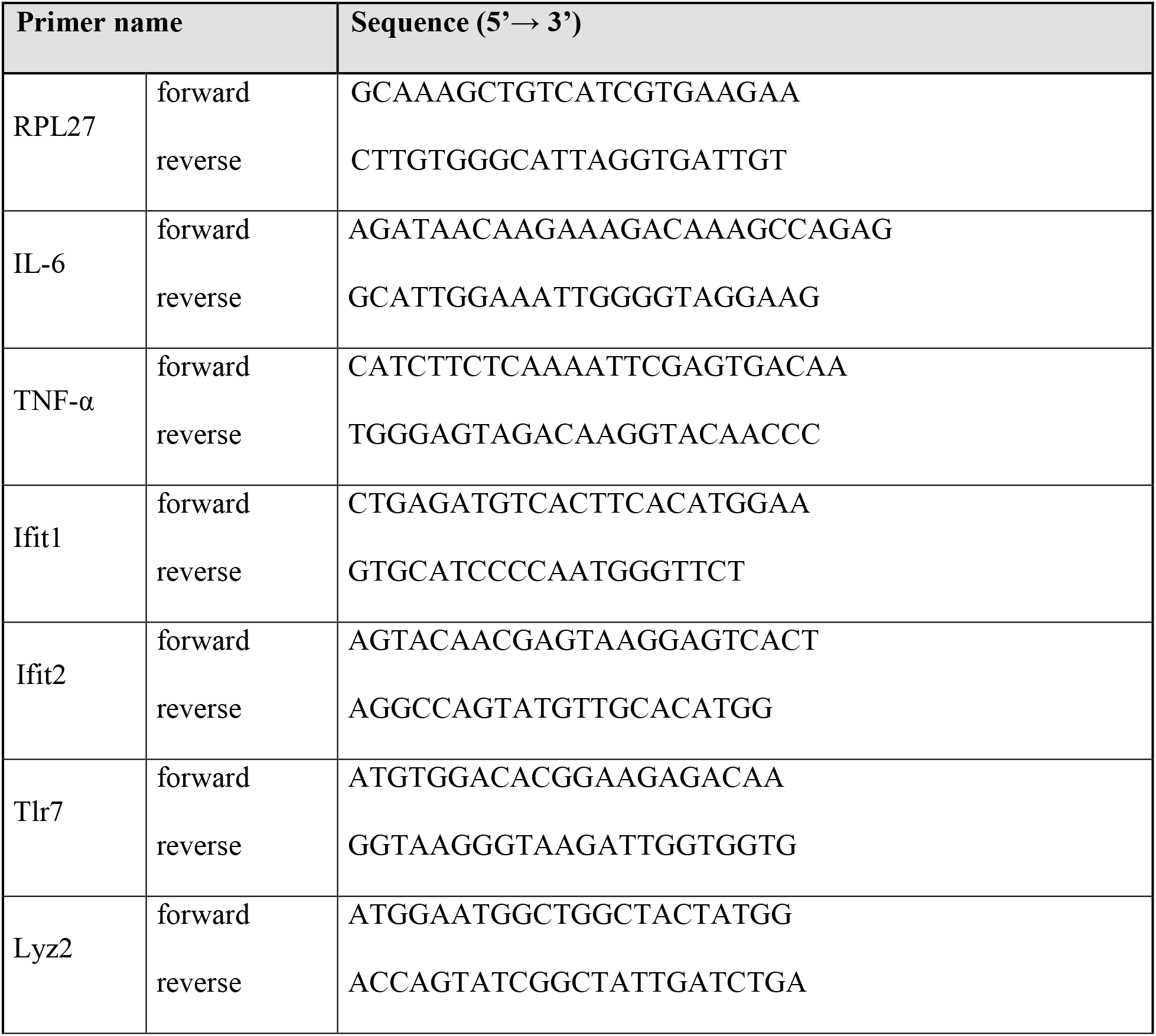
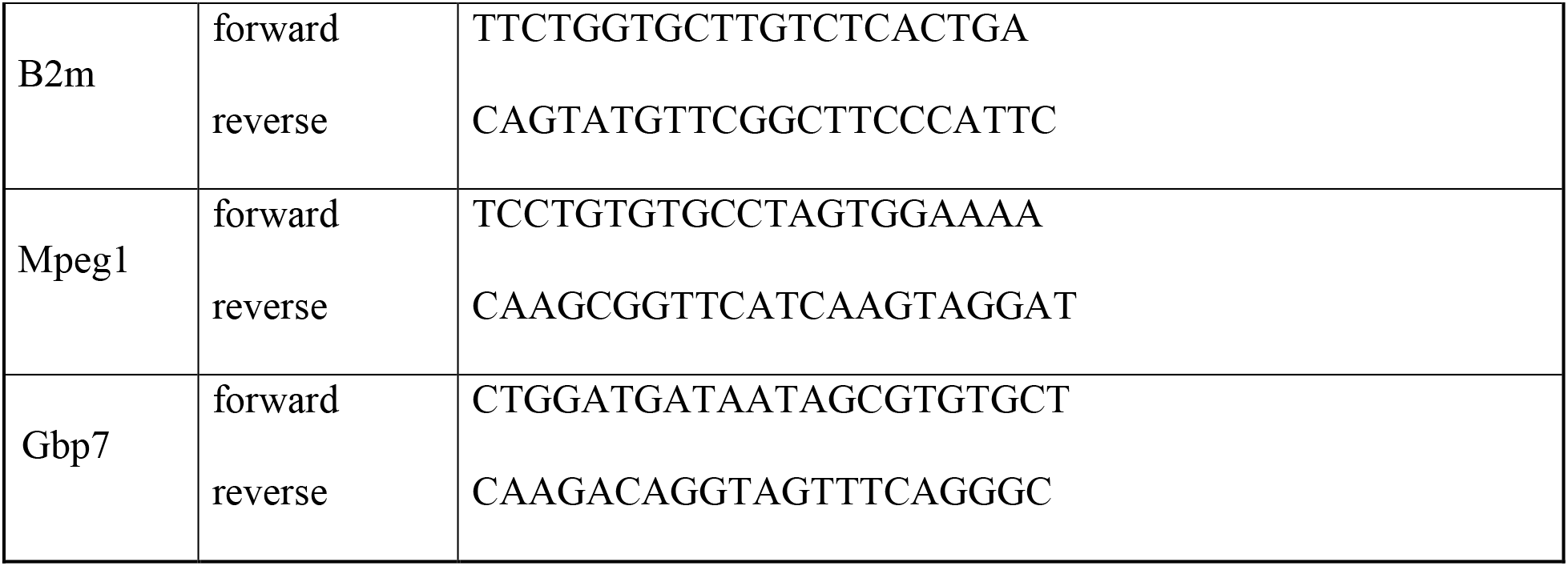
Sequences of primers used for RT-qPCR.

### Statistical analyses

Statistical analyses were performed using Prism version 9.5.1 (GraphPad, La Jolla, CA). A two-way Analysis of Variance (ANOVA) with Sidak’s multiple comparison test was used to determine the significance of the difference in body weight changes. A one-way ANOVA with Tukey’s multiple comparison test was used to analyze viral load and relative gene expression of RT-qPCR data. A p-value of less than 0.05 was considered significant.

## RESULTS

### Loss of body weight is greater in older male C57BL/6J mice infected with SARS-CoV-2 B.1.351 variant

Previous studies have shown that SARS-CoV-2 B.1.351 and other variants with the N501Y mutation in the viral spike protein efficiently infected wild type laboratory mice [21–25]. To determine whether different age groups showed differences in susceptibility to SARS-CoV-2 infection, 4-month, 10-month, and 16-month-old wild type C57BL/6J mice were infected intranasally with 1 x 10^5^ FFU of SARS-CoV-2 B.1.351 variant (South Africa/KRISP-EC-K005321/20204) and monitored for 7 days, with uninfected age/sex-matched mice as controls. No significant loss of body weight was observed in 4-month-old mice infected with SARS-CoV-2 compared to uninfected mice. However, body weight loss was significantly greater in infected, aged male mice at 10-month and 16-month of age, than in female mice of the same age. Loss of body weight peaked at 4 dpi in older male mice (male 6.45 ± 1.15% vs female 0 ± 1.49% in 10-month-old; p <0.01 and male 3.7 ± 1.01% vs female 0.25 ± 1.18% in 16-month-old; p < 0.05; **Fig. 1**), whereas female mice showed no significant loss in body weight compared to uninfected animals. No further loss in body weight was observed in male mice after 4 dpi and all mice survived until the experimental end point of 7 dpi.

**Figure 1.**
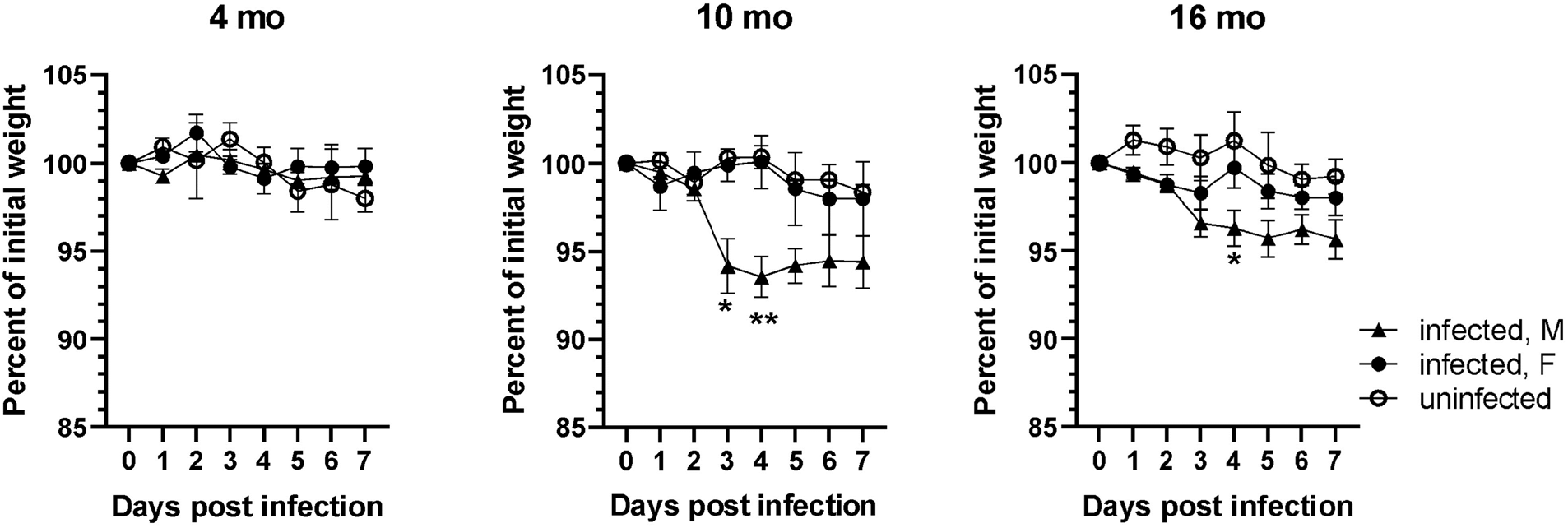
SARS-CoV-2 lineage B.1.135 induced weight loss is higher in older male mice. Male and female C57BL/6J mice at 4, 10, and 16 months of age were infected with 1 x 10^5^ FFU of SARS-CoV-2 lineage B.1.351 via intranasal route. Change in body weight were monitored for 7 days. Error bars indicate standard error of mean (SEM). (n= 3 to 4 in each group). *p <0.05; **p <0.01.

### Older male C57BL/6J mice are more susceptible to SARS-CoV-2 infection than age-matched female mice

The level of viral RNA and infectious virus titer in the lung homogenate of mice were assessed at 7 dpi. As shown in **Fig. 2A**, viral RNA was detectable by RT-qPCR in most of the lungs of SARS-CoV-2 infected mice at 7 dpi. Viral RNA was not detected in the lung of one of the 4-month-old and one of the 16-month-old mice. While there was no significant difference in viral RNA copies between different age groups, there was a trend towards increased viral RNA in the lung in aged, 16-month-old mice compared to 4– and 10-month-old mice (**Fig. 2A**). In addition, the viral RNA load in the lungs of female 4-month and 16-month-old mice was lower than that in male miceat comparable ages, although the difference did not reach statistical significance (4-month p = 0.057; 16-month p = 0.200) (**Fig. 2B**). Infectious viral load in the lung was measured by focus forming assay on Vero E6 cells. Infectious virus was occasionally detected in the lung of younger mice at 7 dpi (**Fig. 2C**) with 14% of 4-month-old (1 out of 7) and 12.5% of 10-month-old (1 out of 8) mice having detectable virus. In contrast, 75% (6 out of 8) of 16-month-old mice harbored detectable levels of infectious virus in the lung at 7 dpi, which was significantly higher than virus in 4– and 10-month-old mice (**Fig. 2C**). Analysis of data based on sex showed that 1 out of 4 male 4-month–old and 1 out of 4 female 10-month-old mice had detectable virus. While all of the aged, 16-month-old male mice had infectious virus in the lung, only 50% (2 out of 4) of female mice showed detectable virus in this age group at 7 dpi (**Fig. 2D**). Collectively, these results indicate that more infectious virus was recovered at 7 dpi in older mice compared to younger mice given the same viral dose. Furthermore, older male C57BL/6J mice are more susceptible to lung infection by SARS-CoV-2 B.1.351 variant than younger mice.

**Figure 2.**
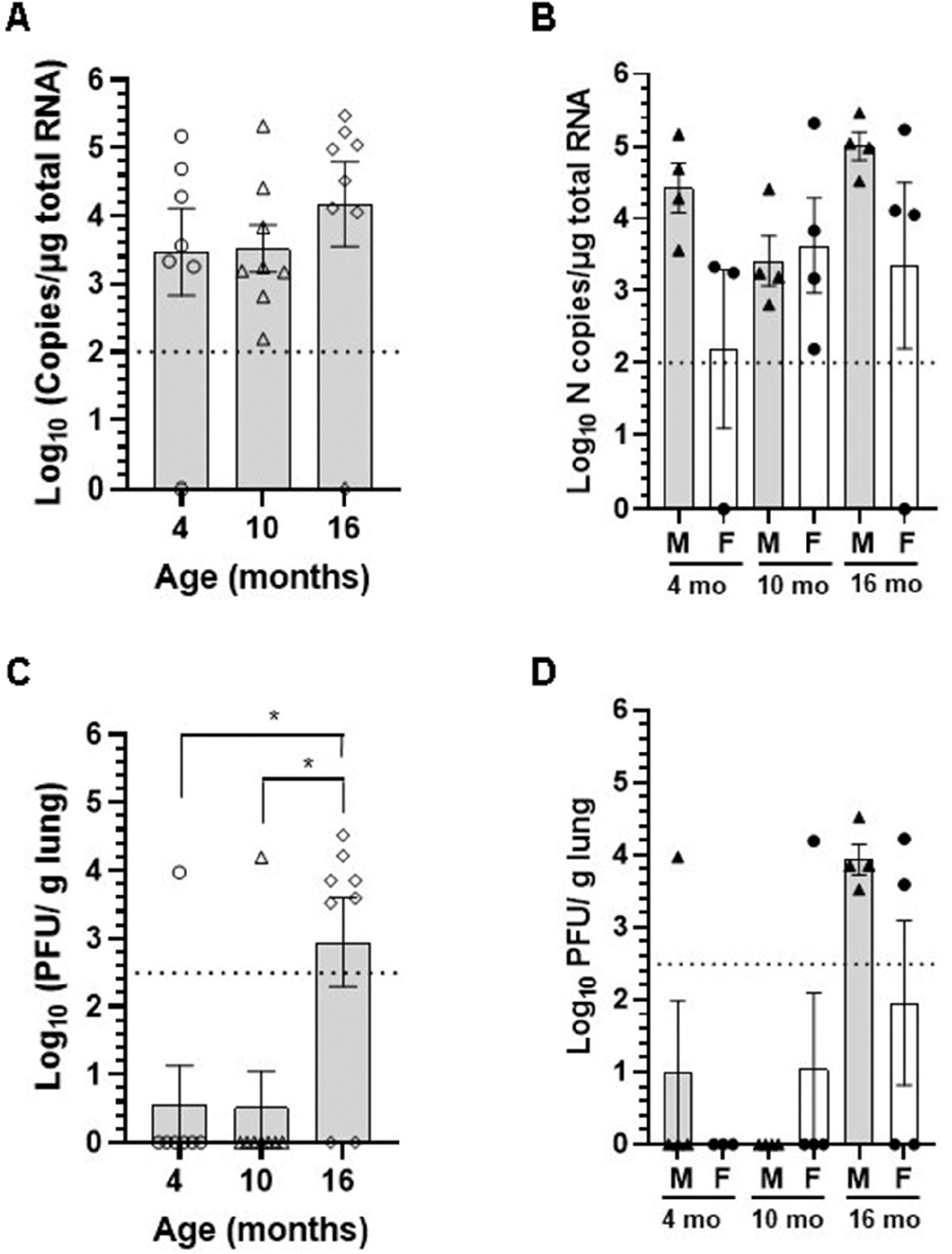
Older male mice are more susceptible to SARS-CoV-2 lung infection. Viral load in the lung homogenate was quantified at 7 dpi by (A and B) RT-qPCR for SARS-CoV-2 N gene and expressed as copies of N gene per μg of total RNA, and (C and D) by focus forming assay on Vero E6 cells. Error bars indicate standard error of mean (SEM). The dotted line indicates the limit of detection of the assay. (n=3 to 4 per group). *p <0.05.

### SARS-CoV-2 infection induces a neuroinflammatory response at 7 dpi despite undetectable virus in the brain

SARS-CoV-2 respiratory infection affects other organ systems including the central nervous system. To investigate the presence of viral RNA in the brain of C57BL/6J mice infected with SARS-CoV-2 B.1.135 variant, RT-qPCR was performed on total brain RNA at 7 dpi. Very low numbers of viral RNA copies (100-150 copies) were detected in two 4-month-old male (2 out of 4), one 10-month-old female (1 out of 4), one 16-month-old male (1 out of 4), and two 16-month-old female (2 out of 4) mice (**Fig. 3A**). Viral RNA was not detectable in brains of majority of mice infected with SARS-CoV-2. Moreover, no infectious virus was detected in the brain of C57BL/6J mice in any of the age groups studied (data not shown). In contrast, we found abundant viral RNA and nucleocapsid protein in brains of 4-month-old K18-hACE2 mice infected with 1 x 10^4^ FFU of SARS-CoV-2 (**Fig. 3B** and **3C**), consistent with previous reports [41–44].

**Figure 3.**
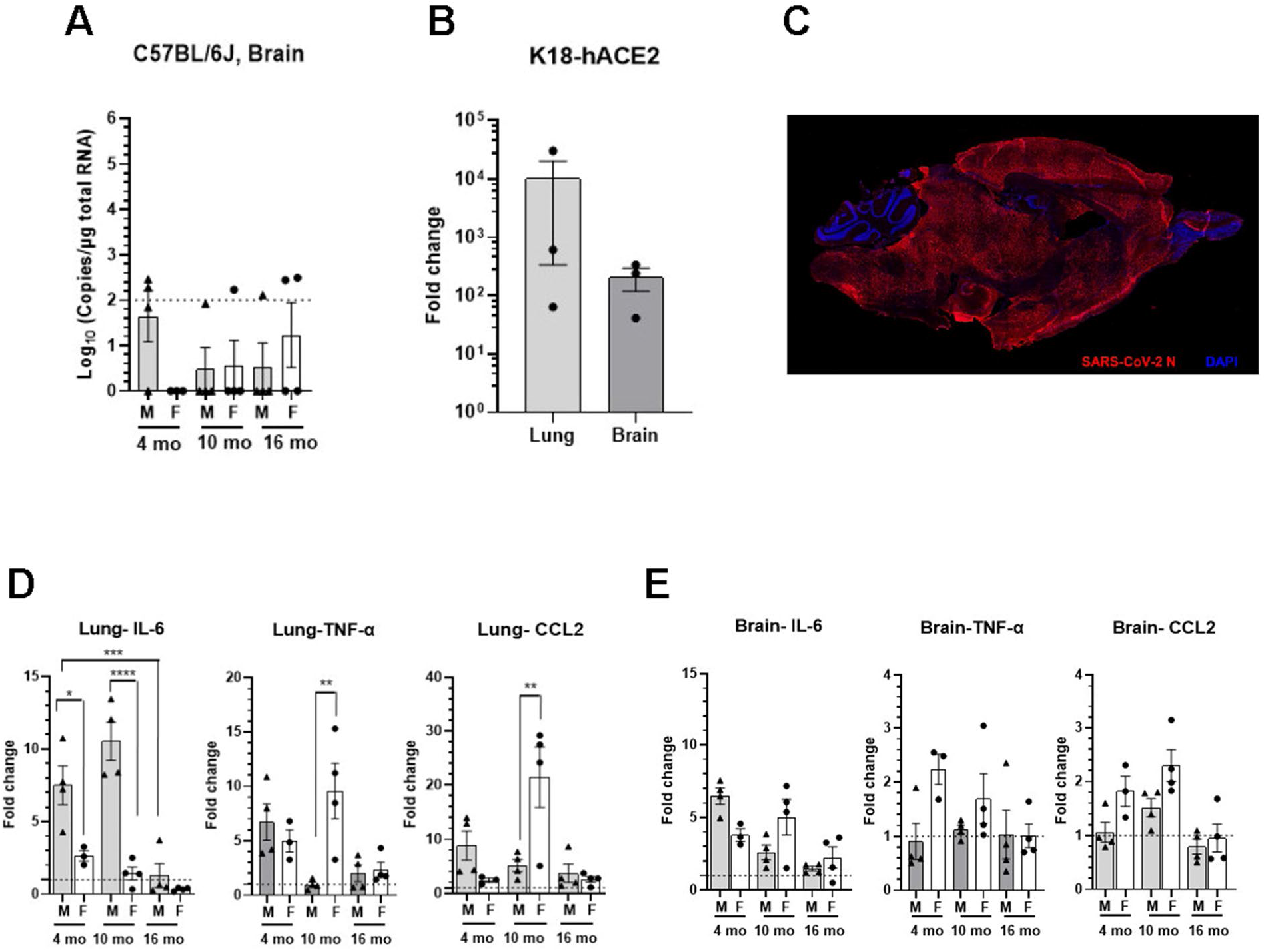
SARS-CoV-2 lineage B.1.135 infection induces an inflammatory response in the brain of C57BL/6J mice despite absence of virus in the brain at 7 dpi. (A) Viral RNA in the brain homogenate of C57BL/6J mice infected with 1 x 10^5^ FFU of SARS-CoV-2 was quantified at 7dpi by RT-qPCR for SARS-CoV-2 N gene and expressed as copies of N gene per μg of total RNA. The dotted line indicates the limit of detection of the assay. (B) Viral RNA in the lung and brain homogenate of K18-hACE2 mice infected with 1 x 10^4^ FFU of SARS-CoV-2 was determined at 7dpi by RT-qPCR for SARS-CoV-2 N gene and expressed as fold change over uninfected control. (C) Representative immunofluorescence image of sagittal brain section of K18-hACE2 mice infected with SARS-CoV-2. Brain sections were immunostained with rabbit polyclonal antibody to SARS-CoV-2 nucleocapsid (N) protein followed by Alexa Fluor 594 conjugated goat anti-rabbit IgG (red). Nucleus were stained with DAPI (blue). (D and E) Relative mRNA expression levels of IL-6, TNF-α, and CCL2 in the (D) lung and (E) brain of C57BL/6J mice infected with SARS-CoV-2 were analyzed by RT-qPCR. Error bars indicate standard error of mean (SEM). The dotted line represents the mean expression level in uninfected control. (n=3 to 4 per group). *p <0.05, **p <0.01, *** p<0.001, ****p <0.0001.

Inflammatory cytokine and chemokine mRNA expression levels at 7 dpi were measured in lungs and brains by RT-qPCR, including IL-6, TNF-α, and CCL2 (**Fig. 3D** and **3E**). As expected, cytokine expresssion levels in the lung were higher in infected animals at all age groups. A significant upregulation of IL-6 mRNA was observed in the lung of 4-month and 10-month-old male mice, compared to age-matched female mice (**Fig. 3D**). In contrast, IL-6 expression was significantly lower in females and notable in males at 16-months of age. TNF-α and CCL2 expression in the lung was higher in infected animals compared to uninfected. However, a trend towards reduced expression of TNF-α and CCL2 was observed in lungs of 16-month-old male mice. In contrast, 10-month-old female mice exibited significatly higher mRNA levels of TNF-α and CCL2 in the lung (**Fig 3D**).

IL-6 expression in the brain was higher in infected animals compared to unifected age matched controls (**Fig. 3E**). Higher expression of IL-6, TNF-α, and CCL2 were observed in 10-month-old female mice compared to male mice, although the difference did not reach statistical significance due to relatively large variations in data (**Fig. 3E**). Elevated levels of TNF-α, and CCL2 were also observed in the brain of 4-month-old female mice. Interestingly, 16-month-old male and female mice showed similar expression levels of IL-6, TNF-α, and CCL2 in the brain post infection.

### SARS-CoV-2 infection induces an innate immune response in the brain

To examine the global effects of SARS-CoV-2 infection on the brain transcriptome, RNA sequencing was performed on brain tissue homogenate collected at 7 dpi from 10-month-old SARS-CoV-2-infected (n = 6) and uninfected C57BL/6J mice (n = 5). Differential gene expression analysis revealed 1067 differentially expressed genes (DEGs). Of these, 478 genes were significantly upregulated (log FC > 1 and adjusted p < 0.05) and 589 genes were significantly downregulated (log FC < 1 and adjusted p < 0.05) in SARS-CoV-2-infected mice (**Fig. 4A**). Hierarchical clustering analysis of significant DEGs (adjusted p-value < 0.05) revealed distinct transcriptional profiles corresponding with SARS-CoV-2 infection (**Fig. 4B**).

**Figure 4.**
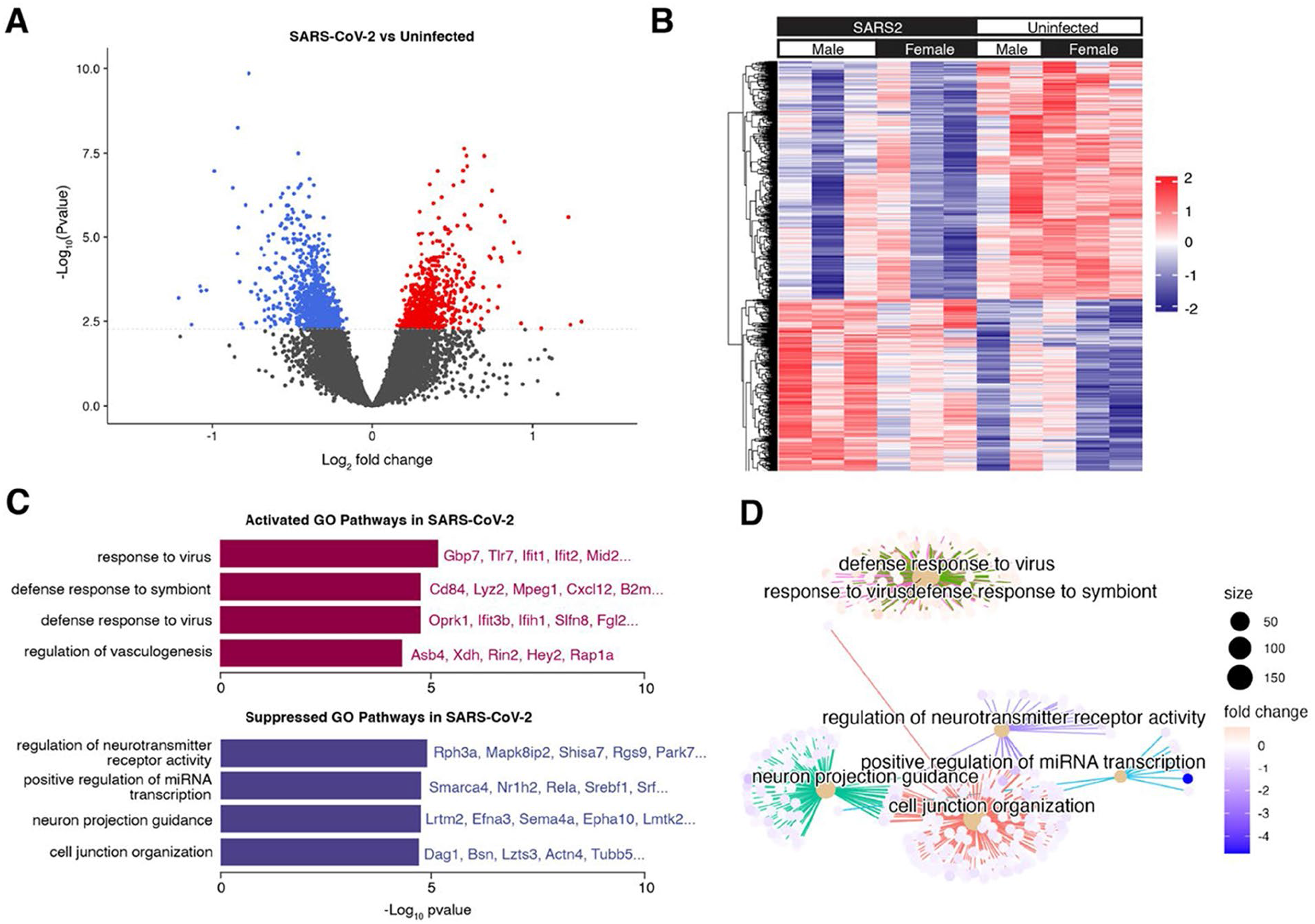
Transcriptional response to SARS-CoV-2 infection. (A) Volcano plot indicating differentially expressed genes (DEGs) (adjusted p < 0.05) of mice infected with SARS-CoV-2 vs. control. Upregulated DEGs in SARS-CoV-2 infected mice are indicated in red, and downregulated DEGs in SARS-CoV-2 infected mice are indicated in blue. The dotted line corresponds to adjusted p = 0.05. (B) Hierarchical clustering heatmap view of DEGs. DEGs are presented in the dendrogram along the Y-axis. Each column contains expression values for an individual mouse, with groups indicated by the color bars along the X-axis. The expression levels from low to high are represented as a color gradient from blue to red, respectively. (C) Barplot visualization of the top 8 (by adjusted p-value) enriched GO biological process pathways. The significance level is indicated as –Log10(adjusted p-value). (D) Cnetplot visualization of the genes in the top 8 (by adjusted p-value) enriched GO biological process pathways. Connections between genes and pathway nodes are color coded. The size of each node represents the number of overlapped genes in each GO term and the color represents log-fold change of each gene between groups.

We performed gene set enrichment analysis (GSEA) to identify the major pathways responsible for the differences between groups (**Table S1**). The top upregulated Gene Ontology (GO) biological process terms in SARS-CoV-2 mice were associated with defense response to virus and other organisms. Among these pathways, common genes included *Tlr7, Ifit1, Ifit2, Ifih1*, and *Gbp7*, all of which play a role in the innate immune response. The top downregulated GO biological process terms in SARS-CoV-2 mice were associated with neuroreceptor activity, axon development, and cell junction organization (**Fig. 4C** and **4D**). Together these results suggest that SARS-CoV-2 may disrupt the blood-brain barrier and promote neuro-axonal injury, in addition to triggering the innate immune response in the brain.

To validate the bulk RNA-seq data, selected immune pathway genes that were upregulated in SARS-CoV-2 infected mice compared to controls (**Fig. S1, Table S2**) were tested by RT-qPCR. The mRNA expression levels of *Ifit1, Ifit2, Tlr7, Lyz2, B2m, Mpeg1*, and *Gbp7* were evaluated in 4-month, 10-month, and 16-month-old mice (**Fig. 5**). Consistent with the RNA-seq data, RT-qPCR results showed significant upregulation of *Ifit1, Lyz2*, and *Mpeg1* mRNA in the brain of 10-month-old SARS-CoV-2 infected mice compared to unifected controls. Although, the expression levels of *Ifit2, Tlr7*, and *B2m* were higher than controls, they did not reach statistical significance. There was no change in the expression of *Gbp7* compared to uninfected controls. Further analysis of gene expression data from different age groups of male and female mice revealed that there was age-dependent decrease in expression of *Ifit1*and *Lyz2* genes in male mice (**Fig. 5**). While no age dependent change was observed for *Lyz2* expression in female mice, a non-significant trend towards higher *Ifit1* expression levels was observed in older female mice. Interestingly, the expression levels of both *Ifit1* and *Lyz2* were greater in 16-month-old female mice compared to age-matched male mice. *Mpeg1* expression was elevated only in 10-month-old female mice. These results demonstrate that age and sex modify the expression of innate immunity-related genes in the brain in response to SARS-CoV-2 infection.

**Figure 5.**
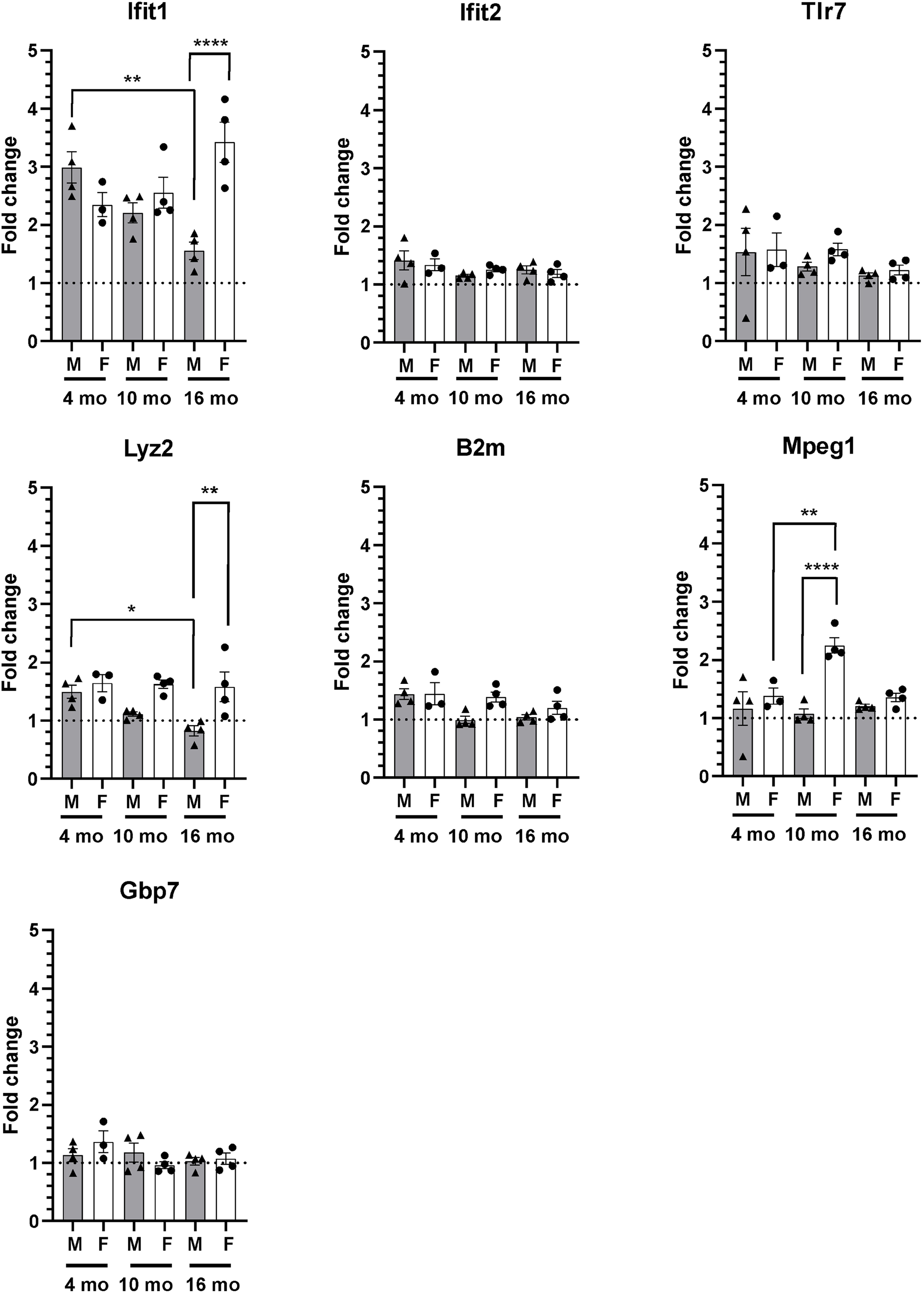
SARS-CoV-2 lineage B.1.135 infection induces an innate immune response in the brain of C57BL/6J mice at 7 dpi. Relative gene expression levels of Ifit1, Ifit2, Tlr7, Lyz2, B2m, Mpeg1, and Gbp7 were analyzed by RT-qPCR. The ΔCT values were normalized to Rpl27 gene expression and represented as fold change (2^-ΔΔCT^) over uninfected control. The dotted line represents the mean expression level in uninfected control. (n=3 to 4 per group). *p <0.5, **p <0.01, ****p <0.0001.

## Discussion

Increased age is considered as one of the major risk factors for severe COVID-19 outcomes [27–29, 45]. Epidemiological studies suggest that patients over 65 years of age account for 80% of COVID-19 hospitalization with 20-fold higher mortality rate compared to those under 65 years [30]. However, comorbidities such as cardiovascular disease, diabetes, and chronic lung diseases also increase with age and may influence the severity of COVID-19 outcome in older patients. Emerging evidence also indicate that COVID-19 severity and mortality are higher among men than women [31–34, 46]. In the present study, we investigated the effect of age on SARS-CoV-2 pathogenesis by comparing the response to infection in 4-, 10-, and 16-months old C57BL/6J mice. Using age/sex-disaggregated data from SARS-CoV-2 infected mice, we assessed the impact of age and sex on SARS-CoV-2 pathogenesis. We used a naturally occurring isolate of SARS-CoV-2 beta variant, B.1.135, capable of infecting wild type laboratory mice to induce severe pathological lesions and inflammatory response in the lung [22–24].

Our data on SARS-CoV-2 infection in a wild-type mouse model reflects the age and sex dependent increase in disease severity reported in humans. We found that the severity of disease outcomes increased in older male mice compared to young or female mice as indicated by the significant loss of body weight at 3 to 4 dpi and increased viral load in the lung at 7 dpi. Corroborating our work, earlier studies reported no significant weight loss in 8-week-old young adult C57BL/6 mice infected with the B.1.135 variant, despite pathological lung lesions and inflammatory responses [21–23]. However, Yasui *et al* reported weight loss in infected 8-week-old Balb/c mice or when infected at higher viral dose (1 x 10^6^ pfu) in C57BL/6 mice [24]. These authors also reported increased weight loss and fatality in aged Balb/c mice [24]. These reports suggest that in addition to age and sex, the strain/genectic background of mice could be an important component of the SARS-CoV2 murine model.

Neurotropism of SARS-CoV-2 is still under debate. Several *in vitro* studies showed that the cells of the CNS, in particular astrocytes and neurons, support SARS-CoV-2 replication [47–53]. Viral RNA was either absent [54–57] or present in low levels in a subset of the autopsy brain samples of fatal COVID-19 patients [5, 53, 58–61]. Consistent with these findings in humans, SARS-CoV-2 RNA was not detectable in the brain of majority of infected wild-type mice, with some brains containing very low levels of viral RNA, while no infectious virus was detected in any of the brains in the present study.

COVID-19 is characterized by pronounced inflammation, excess cytokine production, and alterations in both the innate and adaptive immune responses [62]. Although the underlying mechanisms contributing to brain pathology are not fully understood, we observed significant changes in the transcriptome of SARS-CoV-2 infected brains that point to the involvement of the innate immune system. DEG and GSEA analysis identified upregulation of several genes that modulate the innate immune response. Notable among them is *Tlr7*, a type of toll-like receptor, which is responsible for activation of innate immune responses and regulation of cytokine production [63]. TLR7 was identified as a cellular detector of ssRNA of SARS-CoV-2, which leads to inflammasome activation and production of pro-inflammatory cytokines and interferons via NF-κB [64–67]. Recent reports link genetic *Tlr7* deficiencies with more severe COVID-19 infection in young individuals, connecting TLR7 with host resistance and disease outcome [68–70].

Using a mouse-adapted SARS-CoV-2, Beer *et al* showed that the age-dependent increase in disease severity is due to impaired interferon response in older mice [71]. IFIT1 and IFIT2 (also known as ISG56 and ISG54, respectively) are interferon induced proteins primarily known for their roles in antiviral defense by restricting viral replication and modulating immune responses to combat viral infections [72]. *Ifit1* and *Ifit2* are induced in response to dsRNA, type I and type II IFNs, and infection by various viruses [73] These factors have been shown to inhibit virus replication by binding to eIF3 and limiting translation of viral mRNA [74, 75]. IFIT1 and IFIT2 exert antiviral activity against the SARS-CoV-2 virus by binding directly to capped viral mRNA to inhibit translation or replication [76]. Although we did not assess the level of expression in the lung, *Ifit1* expression was higher in the brain of infected 10-month-old mice compared to controls, implicating IFIT in modulating the host immune response. Furthermore, we found an age-dependent decrease in gene expression of *Ifit1* in the brain of male mice when assessed by RT-qPCR. *Ifit1* expression was lowest in 16-month-old male mice, the group that had maximum viral load in the lung, suggesting impaired *Ifit1* expression in older male mice. Intriguingly, age-dependent reduction in *Ifit1* expression was not observed in female mice.

Furthermore, several genes including *Mpeg1* and *Cd84* are involved in macrophage activation and phagocytosis [77, 78]. Macrophages act as important sentinel cells in peripheral organs where they monitor surrounding tissue for invading pathogens, ingest and kill pathogens, produce and secrete cytokines/chemokines to regulate the immune response [79]. Many recent studies highlight the role of macrophages in the pathogenesis of SARS-CoV-2 virus infection in the lungs [79–82]. However, there remains a gap in the understanding of neuroinflammatory responses associated with COVID-19 in the brain. Taken together, our results imply that macrophages might play a crucial role in the progression of SARS-CoV-2 infection in the brain; however, further investigation is required to elucidate the exact factors driving macrophage activation and underlying molecular events governing their phenotype at various stages of infection.

Pathways related to cell junction organization and regulation of vasculogenesis were altered in SARS-CoV-2-infected brains, as seen in the GO enrichment analysis. Several proteins involved in maintaining blood-brain barrier (BBB) integrity were downregulated, including Cldn5, a member of the claudin family. Claudins are key components of tight junctions between adjacent endothelial cells that regulate the permeability of the BBB [83, 84]. Multiple proteins with functions related to regulation of endothelial cell-matrix adhesion or cell-cell junction organization (Ajm and Pard6a, Nectin1, Rcc2, Cbln1) were also expressed at significantly lower levels in infected brains, suggesting that SARS-CoV2 infection led to disruption of BBB integrity and neurovascular function.

Consistent with the GO analysis observations, we also found a significant decrease in expression of cadherin proteins (Cdh1, Cdh22, Cdh15, Cdh19) in SARS-CoV-2 infected brains. Cadherins are transmembrane proteins that mediate cell–cell adhesion and cell-cell junction organization [85, 86]. These findings furthur support the possiblility of BBB disruption and subsequent activation of the innate immune response in the SARS-CoV-2 infected mice brains. The inflammatory response induced in the brain is likely a result of peripheral immune cells infiltrating the CNS through a compromised BBB, consequently contributing to neuroinflammation and neuronal cell death, as indicated by suppression of axonogenesis and down regulation of neurotransmitter receptor activity pathways in infected brain.

The results of the present study are consistent with previous *in vitro* experiments that demonstrate that SARS-CoV-2 disrupts the BBB [87–90]. We have shown transcriptional *in vivo* evidence that cerebrovascular changes and inflammation occur upon infection, which provide insights into the mechanisms underlying the neurological symptoms of COVID-19. A comprehensive investigation into mechanisms by which SARS-CoV-2 induces BBB disruption and immune dysfunction may offer novel avenues for therapeutic interventions in the management of COVID-19.

## Conclusions

In summary, our findings indicate that SARS-CoV-2 variant B.1.135 infection induces a neuroinflammatory response despite the lack of detectable virus in the brain. Age and sex modify the susceptibility and severity of SARS-CoV-2 infection. An activated innate immune response, compromised BBB integrity, and supressed neuronal activities underlie the pathogenic impact of SARS-CoV-2 infection on the brain. Further studies are required to determine the long-term neuropathological changes due to SARS-CoV2 infection and to elucidate the underlying mechanisms that drive the neurological manifestaions of COVID-19.

## Declarations

### Ethics approval

All studies using infectious SARS-CoV-2 were conducted in certified BSL-3/ABSL-3 facilities at the University of Minnesota (UMN). The protocols and procedures used in the following studies have been approved by the UMN Institutional Biosafety Committee. Animal studies were conducted in ABSL3 facilities and managed in an investigator managed housing area as per protocols approved by the UMN Institutional Animal Care and Use Committee and in accordance with the Guide for the Care and Use of Laboratory Animals from the National Institute of Health.

### Availability of data and Materials

The datasets used and/or analysed during the current study are available from the corresponding author on reasonable request. The RNAseq data are available in the Gene Expression Omnibus repository (GEO Series accession number GSE237092).

## Abbreviations

ACE 2: Angiotensin-converting enzyme 2
ANOVA: Analysis of variance
B2M: Beta-2-microglobulin
BBB: Blood-brain barrier
CCL2: Chemokine ligand 2
CD84: Cluster of differentiation 84
CNS: Central nervous system
COVID-19: Coronavirus disease 2019
DGE: Differential gene expression
dsRNA: Double-stranded ribonucleic acid
GBP7: Guanylate binding protein 7
GO: Gene ontology
GSEA: Gene set enrichment analysis
IFIT: Interferon induced protein with tertratricopeptide repeats
IFN: Interferon
IL-6: Interleukin 6
ISG: Interferon-stimulated gene
Lyz2: Lysozyme 2
MPEG1: Macrophage expressed 1
RT-qPCR: Real-time quantitative polymerase chain reaction
SARS-CoV-2: Severe Acute Respiratory Syndrome Coronavirus 2
SEM: Standard error of mean
TJ: Tight junction
TLR7: Toll like receptor 7
TNF-α: Tumor necrosis factor α

## Acknowledgements

The authors thank the UMN BLS-3 management team, including Thien Sam, Jordan Merhar, Aubree Kees, and Paige Horsch for their assistance in high containment work. We thank Dr. Declan Schroeder, Veterinary Population Medicine department, College of Veterinary Medicine, University of Minnesota, for verifying the virus isolate by sequencing. We also acknowlege the technical help from Swathi Radha and Andrea Gram for experiments described in this manuscript. We thank the University of Minnesota Informatics Institute and Juan E. Abrahante Lloréns, and Ying Zhang for assisting with RNA-seq data analysis.

## Author Contributions

VK conducted the experiments, curated the data, and conducted a formal analysis. AC conducted the RNA-seq data analysis. VK, AC, HK, and SV worked on the methodology and wrote the first draft of the manuscript. LL, WL, and MC conceived the study, supervised the progress of all experiments, interpreted the results, and edited and finalized the manuscript. All authors reviewed and approved the final manuscript.

