## Supplementary material for "Impact of age and sex on neuroinflammation following SARS-CoV-2 infection in a murine model": Fig. S1, Table S2

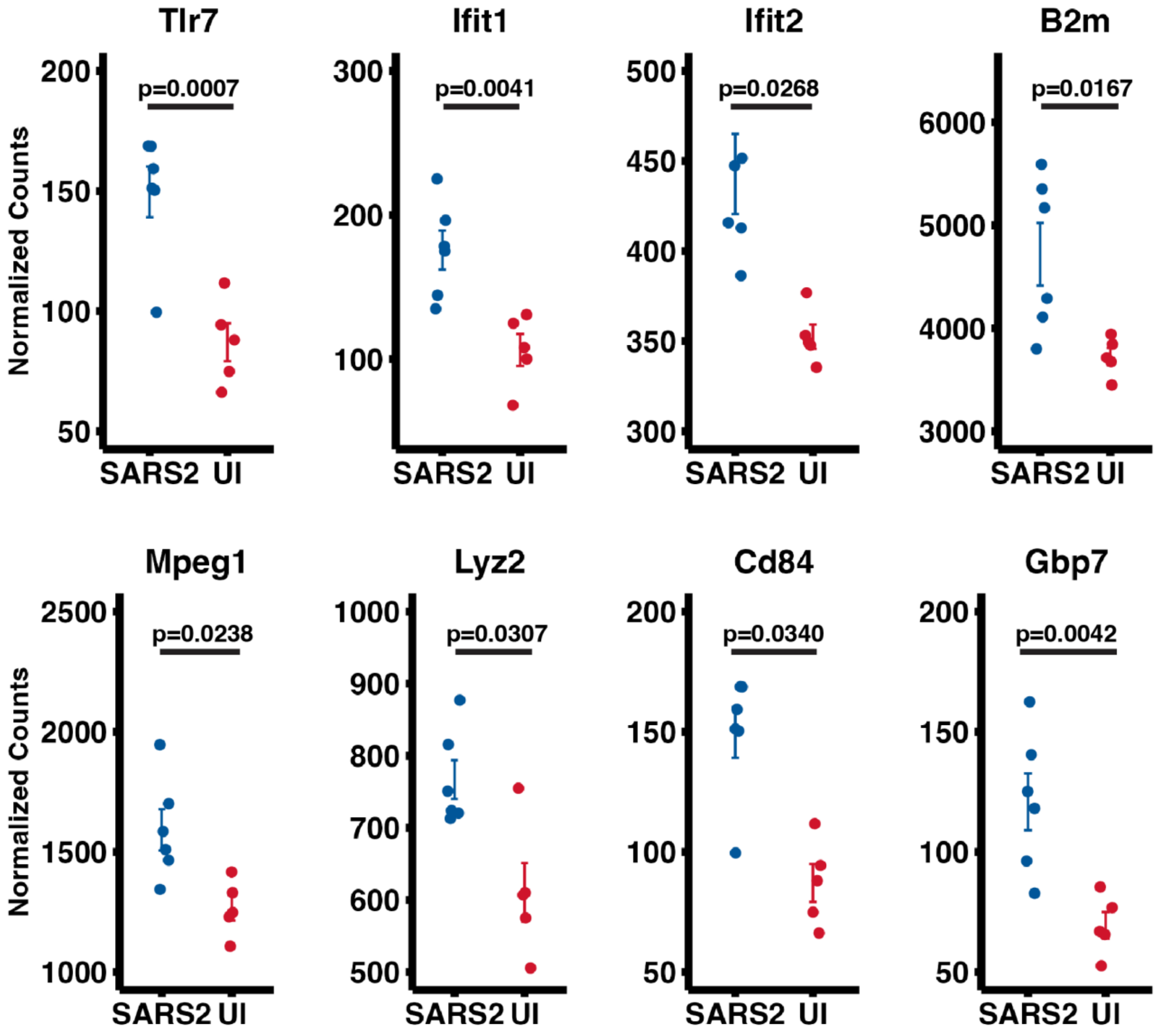


**Fig S1.** Plots showing DESeq2 normalized gene expression level of selected DEGs shared among GO immune pathways upregulated in SARS-CoV-2 mice compared to uninfected mice. Benjamini-Hochberg corrected p values are based on DEseq2 analysis. Mean values ± SEMs are shown.

**Table S1.** Gene Set Enrichment Analysis (GSEA) enriched GO pathways.

| ***Pathway*** | ***Size*** | ***ES*** | ***NES*** | ***pvalue*** | ***FDR*** |
|---|---|---|---|---|---|
| *regulation of vasculogenesis* | 16 | 0.75 | 2.18 | 0.00 | 0.026 |
| *positive regulation of miRNA transcription* | 42 | -0.57 | -2.11 | 0.00 | 0.026 |
| *bone morphogenesis* | 79 | -0.49 | -2.10 | 0.00 | 0.026 |
| *neuron projection guidance* | 214 | -0.36 | -1.80 | 0.00 | 0.026 |
| *defense response to virus* | 197 | 0.37 | 1.79 | 0.00 | 0.026 |
| *defense response to symbiont* | 197 | 0.37 | 1.79 | 0.00 | 0.026 |
| *axon guidance* | 213 | -0.36 | -1.79 | 0.00 | 0.026 |
| *response to virus* | 236 | 0.35 | 1.75 | 0.00 | 0.026 |
| *cell junction organization* | 671 | -0.26 | -1.49 | 0.00 | 0.028 |
| *regulation of neurotransmitter receptor activity* | 60 | -0.53 | -2.14 | 0.00 | 0.028 |
| *miRNA transcription* | 56 | -0.53 | -2.09 | 0.00 | 0.028 |
| *axon development* | 474 | -0.29 | -1.58 | 0.00 | 0.028 |
| *axonogenesis* | 431 | -0.29 | -1.60 | 0.00 | 0.030 |
| *defense response to other organism* | 595 | 0.26 | 1.47 | 0.00 | 0.030 |

**Table S2.** Cell type expression and function of selected immune-pathway genes

| Gene Name | Cell Type | Function | References |
|---|---|---|---|
| **Tlr7** | Macrophage, Microglia, astrocytes | Innate immune receptor that recognizes single-stranded RNA | (Li et al., 2021; Michaelis et al., 2019) |
| **Ifit1and Ifit2** | Low basal expression, enhanced in response to IFNs and viral infection | Binds to capped RNA to inhibit viral replication and translational initiation | (Mears and Sweeney, 2020; Pidugu et al., 2019; Terenzi et al., 2007) |
| **B2m** | Distinct expression in macrophages and immune cells | Participate in formation of MHC class I complexes, facilitates presentation of viral antigens on infected cells to T cells for immune recognition | (Wang et al., 2022) |
| **Mpeg1** | Most abundant in immune cells | Promotes phagocytosis and antimicrobial activity in macrophages | (Bayly-Jones et al., 2020) |
| **Lyz2** | Myeloblasts, macrophages, and neutrophils | Exhibit antimicrobial activity by catalyzing the hydrolysis of peptidoglycan in bacterial cell walls | (Orthgiess et al., 2016) |
| **Cd84** | Monocytes, macrophages, granulocytes, and dendritic cells | Signaling molecule that regulates immune cell activation and cell-cell interactions | (Cuenca et al., 2019; Sintes et al., 2010) |
| **Gbp7** | B-cells, T-cells, NK-cells, monocytes, granulocytes and dendritic cells | Interferon (IFN)-inducible GTPase that plays important roles in host defense against a range of bacterial, viral and protozoan pathogens | (Steffens et al., 2020) |
